# Posttraumatic Stress Disorder and the Social Brain: Affect-Related Disruption of the Default and Mirror Networks

**DOI:** 10.1101/527176

**Authors:** Kevin M. Tan, Lisa J. Burklund, Michelle G. Craske, Matthew D. Lieberman

## Abstract

**Background:** Social impairments, specifically in mentalizing and emotion recognition, are common and debilitating symptoms of posttraumatic stress disorder (PTSD). Despite this, little is known about the neural underpinnings of these impairments, as there have been no published neuroimaging investigations of social inference in PTSD.

**Methods:** Trauma-exposed veterans with and without PTSD (*N* = 20 each) performed the Why/How social inference task during functional magnetic resonance imaging (fMRI). The PTSD group had two fMRI sessions, between which they underwent affect labeling training. We probed the primary networks of the “social brain”—the default mode network (DMN) and mirror neuron system (MNS)—by examining neural activity evoked by mentalizing and action identification prompts, which were paired with emotional and non-emotional targets.

**Results:** Hyperactivation to emotional stimuli differentiated PTSD patients from controls, correlated with symptom severity, and predicted training outcomes. Critically, these effects were generally non-significant for non-emotional stimuli. PTSD-related effects were widely distributed throughout DMN and MNS. Effects were strongest in regions associated with the dorsal attention, ventral attention, and frontoparietal control networks. Unexpectedly, effects were non-significant in core affect regions.

**Conclusions:** The array of social cognitive processes subserved by DMN and MNS may be inordinately selective for emotional stimuli in PTSD. This selectivity may be tightly linked with attentional processes, as effects were strongest in attention-related regions. Putatively, we propose an attentional account of social inference dysfunction in PTSD, in which affective attentional biases drive widespread affect-selectivity throughout the social brain. This account aligns with numerous findings of affect-biased attentional processing in PTSD.

## Introduction

PTSD is characterized by intrusive trauma-related cognition (e.g. thoughts, dreams, and flashbacks), exaggerated affective responses (e.g. chronic fear, anxiety, and hyperarousal), and—conversely—affective blunting (e.g. anhedonia and emotional numbing) (1). Though PTSD is most commonly associated with affective dysfunction, social dysfunction is ubiquitous and often debilitating in PTSD (2–4). This has led many researchers to emphasize social and interpersonal factors in the development of PTSD and related adversity (5–7). A strong body of behavioral evidence links PTSD with deficits in emotion recognition (8–13) and mentalizing (13–17), a pattern of social cognitive impairment that is distinct from other anxiety disorders (18). Here, emotion recognition refers to perceiving and identifying others’ emotions, while mentalizing refers to reasoning about others’ mental states (e.g. beliefs, desires, and intentions). Emotion recognition can be considered a type of mentalizing, and both are facets of social inference and theory of mind (19–21). Taken together, there is converging behavioral evidence that social inference impairments are common and debilitating symptoms of PTSD. However, little is known about the neural underpinnings of these impairments.

In healthy populations, neuroimaging investigations have revealed that social inference is primarily subserved by two dissociable large-scale neural networks: the mirror neuron system (MNS) and default mode network (DMN) (22–24). MNS is associated with action identification, while DMN is associated with mentalizing (25–28). Mirror neurons, first discovered in macaque frontoparietal cortex, fire when actions are either performed or observed (29–31). In humans, similar sensorimotor “mirroring” responses may have been found in posterior inferior frontal gyrus (pIFG), dorsal premotor cortex (dPMC), inferior parietal sulcus (IPS), and lateral occipitotemporal cortex (LOTC) (22, 32). Such MNS regions have been shown to encode facial expressions (33), body language (34), and other goal-directed motor actions (35–37). During social inference, MNS is thought to represent observable sensorimotor actions (e.g. reaching for a cup) that are used by DMN to infer unobservable mental states and traits (e.g. thirsty) (27, 38, 39). Concordantly, MNS-based sensorimotor encoding appears to precede DMN-based mentalizing (40–43).

DMN regions are consistently recruited by mentalizing (44) and other functions that involve abstract mental state reasoning, such as theory of mind (45), emotion recognition (40), empathy (46), moral cognition (47), social working memory (48), and introspection (49). Though the anatomical distribution of these functions can differ, they often include the core DMN hubs of medial prefrontal cortex (mPFC), posterior cingulate cortex (PCC), and temporoparietal junction (TPJ) (24, 50–53). Aside from social and emotional functions, DMN is broadly associated with internally-oriented cognition (54). However, much of DMN activity occurs during rest, as DMN activation and connectivity are quickly engaged and sustained during the absence of goal-directed cognition (55–58). As such, DMN is widely thought to subserve the “default mode” of mammalian brain function (23, 58, 59).

As detailed above, the neural substrates of social inference are fairly well-characterized in healthy populations. In contrast, little is known about the neurobiology of social inference impairments in PTSD, as there are currently no published neuroimaging studies that directly investigate social inference in PTSD. However, PTSD-related alterations in DMN activity have been found in other social tasks (60), such as script-driven social-emotional imagery (61–63), self-reference (64, 65), self-other reference (66), and face perception (67, 68). Moreover, PTSD-related differences in DMN connectivity are consistently found in resting-state studies (69–73). We are unaware of any reports of PTSD-related effects in regions explicitly defined as MNS. However, MNS appears to overlap substantially with the dorsal and ventral attention networks (74, 75, 32, 58), which are strongly implicated in PTSD-related attentional biases (76–86).

Here, we perform the first neuroimaging investigation of social inference in PTSD. We probed activity in DMN and MNS regions due to their importance in social cognition and PTSD. To this end, we used functional magnetic resonance imaging (fMRI) to record brain activity during the Why/How social inference task (87) in trauma-exposed veterans with and without PTSD. The Why/How task contains mentalizing (*Why*) and action identification (*How*) prompts (Figure 1), which robustly dissociate DMN and MNS activity (Figure 2) (26, 28, 29, 41, 42, 89). We explored whether DMN and MNS responses differentiate PTSD patients from controls, correlate with symptom severity, and predict training outcomes.

**Figure 1.**
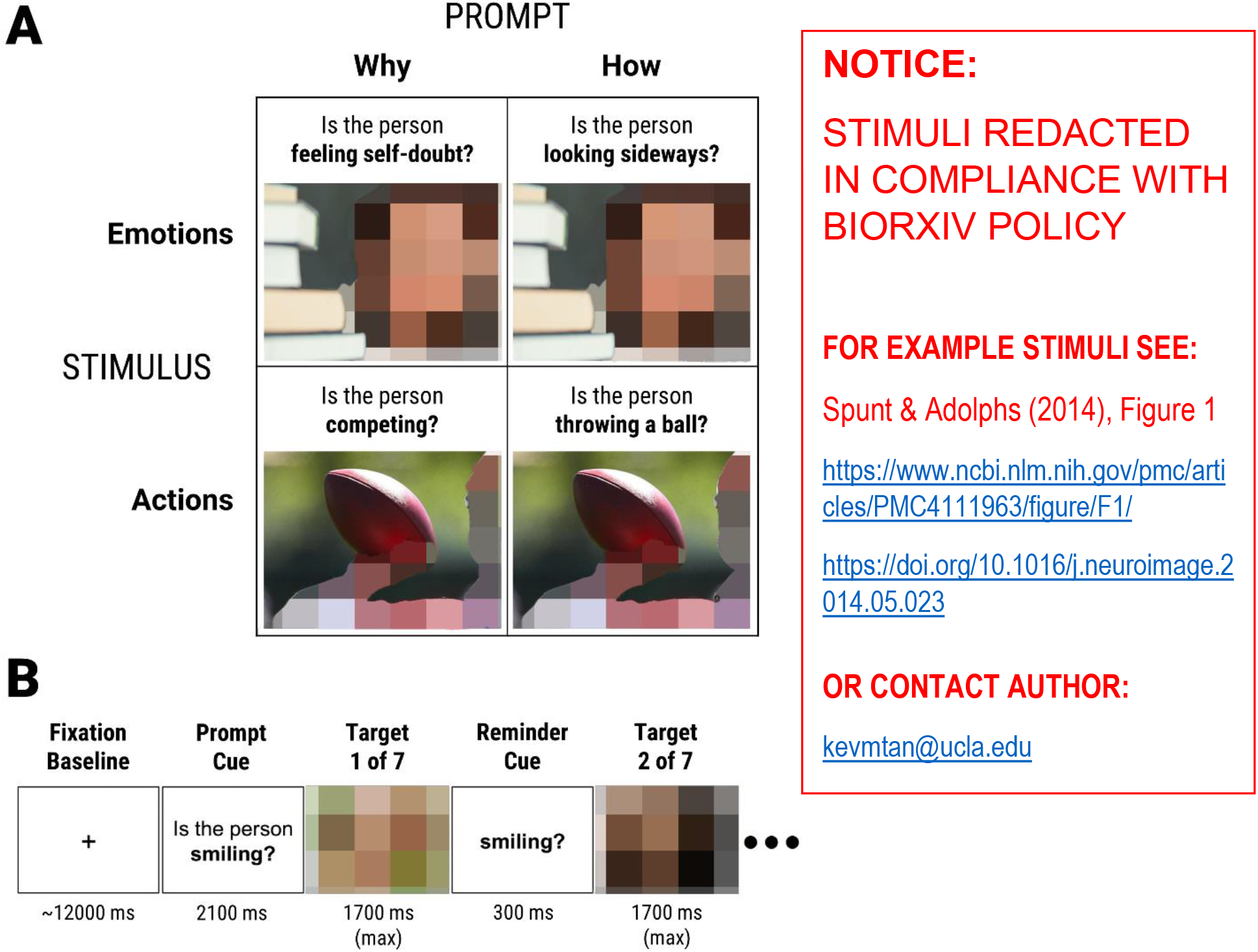
Summary of the standardized Why/How social inference task (Study 3 of Spunt & Adolphs, 2014). **(A)** Diagram of the task’s 2×2 design across Prompt and Stimulus. Each stimulus was shown twice: once with a mentalizing prompt (*Why*), and once with an action identification prompt (*How*). There were two types of stimuli: emotional facial expressions (*Emotions*), and intentional hand actions (*Actions*). **(B)** Sequence of events in a task block. Each block began with a prompt followed by seven target stimuli paired with that prompt. Participants were instructed to make true/false responses as quickly and accurately as possible during the presentation of target stimuli. Target stimuli were presented for 1700 ms or until a response was made. A reminder prompt was shown for 300 ms between target stimuli, and each block was preceded by a fixation baseline period.

## Materials and Methods

### Participants and Procedure

Forty trauma-exposed military veterans were recruited from the Los Angeles area. All participants were exposed to combat trauma, mostly in Iraq and Afghanistan. Twenty participants met DSM-5 criteria for PTSD or other trauma-related disorder, while the control group (*N* = 20) had no current or lifetime psychiatric diagnoses. Diagnostic status was determined by the Clinician-Administered PTSD Scale (CAPS; 90), which was administered by certified research staff. Participants were 18-45 years old, English-speaking, right-handed, and were excluded for serious medical conditions, moderate-to-severe substance abuse, recent changes to medication/psychotherapy, chronic childhood abuse/neglect, and standard fMRI contraindications (e.g. metallic implants, claustrophobia, pregnancy). Participants provided informed consent, and the study was approved by the University of California, Los Angeles (UCLA) institutional review board.

All participants performed baseline pre-treatment assessments involving a clinical interview, questionnaires, and an fMRI scan. Only the PTSD group continued with three weeks of twice-weekly affect labeling training, followed by post-training assessments similar to the pretraining assessments. Affect labeling training involved repeated practice with several computer-based tasks that were designed to strengthen inhibitory capacity (90–93). This training regimen was investigated as proof-of-concept for a novel, brief computerized intervention for PTSD; full methods and data for affect labeling training will be presented in a separate manuscript (94).

In the baseline session, data from 18 PTSD and 17 control participants were used, while the rest were unusable due to lack of task data (*N* = 3), a previous brain tumor (*N* = 1), and non-compliance with fMRI instructions (*N* = 1). Due to participant dropout, only 13 PTSD participants completed the post-training session, and only data from 11 PTSD participants were used, while the rest were unusable due to lack of task data (*N* = 1), a previous brain tumor (*N* = 1), and suspected cannabis intoxication (*N* = 1).

### Why/How Social Inference Task

Participants completed the “fast” version of the standardized Why/How social inference task (95), which corresponds to Study 3 in (87). The task features a 2×2 design across Prompt_[Why, How]_ and Stimulus_[Emotions, Actions]_ (Figure 1A). Stimuli were photographs of emotional facial expressions (*Emotions*) or intentional hand actions (*Actions*). Each stimulus was shown twice, once with each prompt type. *Why* prompts involve mentalizing (e.g. “Is the person competing?), while *How* prompts involve action identification (e.g. “Is the person throwing a ball?”). Thus, there were four conditions: *WhyEmotions, WhyActions, HowEmotions,* and *HowActions*. The task was organized into 16 blocks, four blocks per condition. Blocks began with a text prompt followed by seven target stimuli paired with that prompt (Figure 1B). Participants were instructed to judge whether a prompt was true or false for a target stimulus as quickly and accurately as possible. The task was implemented in PsychophysicsToolbox 3 (96) running on MATLAB 2007a (97). The task was shown in the fMRI scanner via virtual reality goggles at 800×600 resolution, and responses were made through a button box held with the right hand.

### fMRI Acquisition and Preprocessing

fMRI data were acquired at the UCLA Staglin Center for Cognitive Neuroscience using a Siemens TimTrio 3-Tesla MRI scanner. Functional data were collected through T2*-weighted echo-planar image volumes (slice thickness = 3 mm, gap = 1 mm, 36 slices, TR = 2000 ms, TE = 25 ms, flip angle = 90°, matrix = 64×64, FOV = 200 mm). Two structural scans were acquired: a matched-bandwidth T2-weighted anatomical scan (MBW; slice thickness = 4 mm, no gap, 34 slices, TR = 5000 ms, TE = 34 ms, flip angle = 90°, matrix = 128×128, FOV = 196 mm), and a T1-weighted, magnetization-prepared, rapid-acquisition, gradient-echo scan (MPRAGE; slice thickness = 1 mm, gap = .5 mm, 160 slices, TR = 1900 ms, TE = 3.43 ms, flip angle = 9°, matrix = 256 × 256, FOV = 256 mm).

fMRI data were preprocessed via SPM12 (98) and the DARTEL pipeline (99). For each subject and session, functional images were realigned and resliced to the mean functional image to correct for head motion. Then, the MBW was coregistered to the mean functional image, then the MPRAGE was coregistered to the MBW. Afterwards, the MPRAGE was segmented and bias-corrected. The resulting MPRAGE images and segmentation parameters were used to create a sample-specific image template, which was subsequently affine-registered into Montreal Neurological Institute (MNI) space (100). Deformation fields generated in the previous step were used to normalize all images into MNI space, with functional images undergoing integrated spatial smoothing (8 mm, Gaussian kernel, full-width at half-maximum).

### Single-Subject fMRI Analysis

To estimate neural responses to the Why/How task within each participant and session, task timings were specified in SPM12’s general linear model and convolved with the canonical double-gamma hemodynamic response function. Realignment parameters were used as nuisance regressors to dampen the impact of remaining motion artifacts. Data were high-pass filtered at 1/128 Hz to correct for signal drift. Parameter estimates from Why-How contrasts for both stimulus types (WhyEmotions-HowEmotions and WhyActions-HowActions) were used for all subsequent group-level fMRI analyses.

### Group-Level Analyses

Unless otherwise noted, group-level statistical analyses were performed via Matlab 2016b Statistics and Machine Learning Toolbox (101), with linear mixed-effects models (LMEMs) used for hypothesis testing. LMEMs were specified with the maximal random (within-subject) effects structure justified by each analysis, as this has been shown to be ideal for hypothesis testing (102, 103). *Post-hoc* simple effects tests were performed for LMEMs with significant interaction effects. To obtain canonical “main effects,” effects coding was used in multi-factor models with at least one categorical factor, otherwise dummy coding was used (104).

#### Behavioral Analyses

In the Why/How task, pre-training group differences in response time and accuracy were analyzed via standard and logistic LMEMs, respectively. Both LMEMs featured a full-factorial design between Group_[PTSD, Control]_, Prompt_[Why, How]_, and Stimulus_[Emotions, Actions]_. The intercept, Prompt, Stimulus, and Prompt x Stimulus were nested within Subject.

#### Region of Interest (ROI) fMRI Analyses

To interrogate brain regions that subserve social inference, masks of *a priori* ROIs (Figure 2) were functionally defined by the Why-How contrast in an independent dataset featuring healthy participants (*N = 50*; studies 1 and 3 in (87)). A one-sample two-tailed t-test was used to reveal brain regions that are differentially modulated by the *Why* and *How* conditions. The whole-network DMN (*Why* > *How*) and MNS (*How* > *Why*) masks (Figure 2A) were defined with a threshold of *p* < 0.001. A more stringent threshold of *p* < 1×10^−6^ was used to define ROIs within DMN and MNS (Figure 2B) that are thought to be key nodes of each network (75, 105).

**Figure 2.**
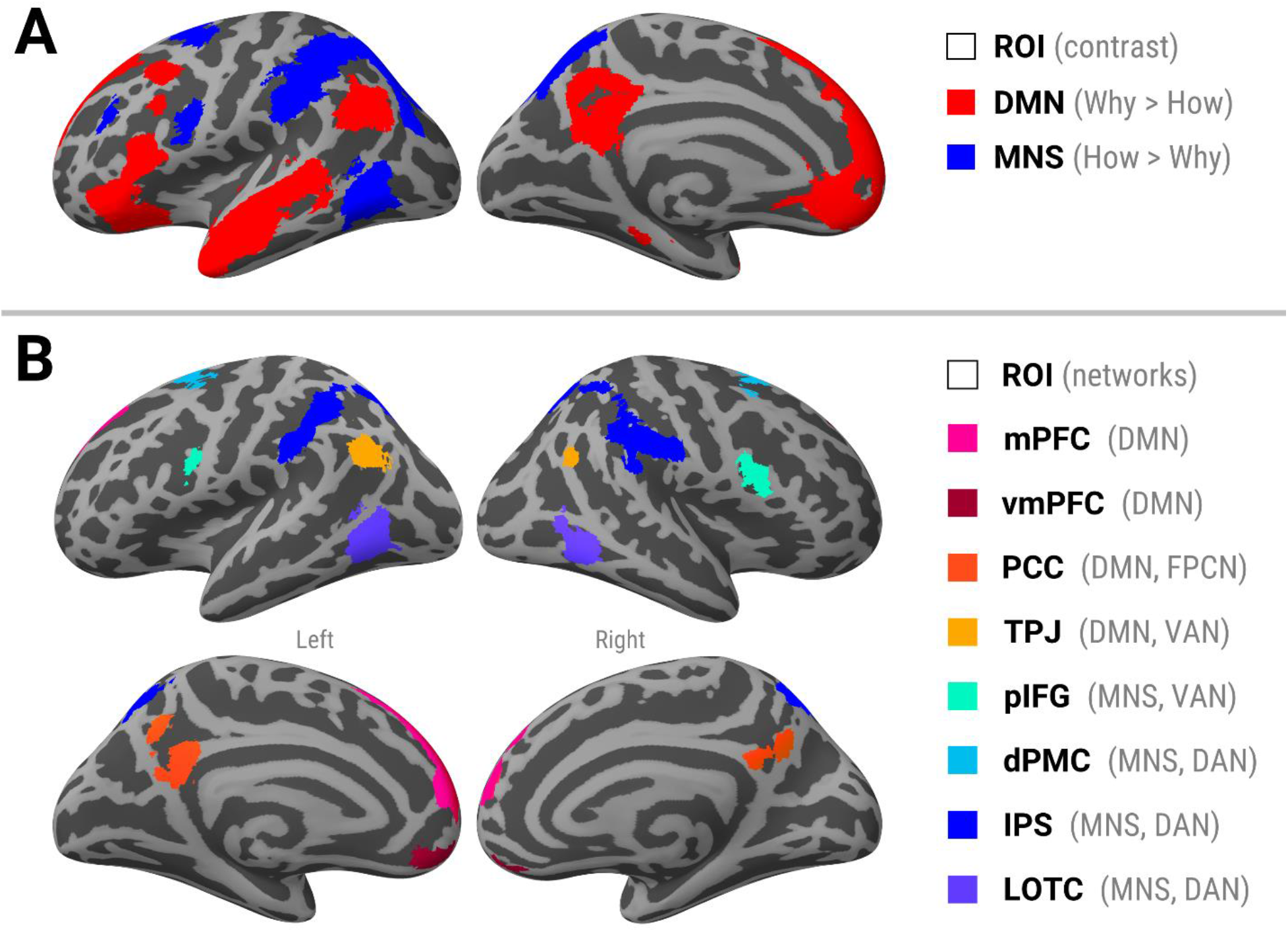
*A priori* ROI masks defined by the Why-How contrast in an independent dataset (Spunt & Adolphs, 2014). The Why-How contrast dissociates DMN and MNS regions that are selective for mentalizing and action identification, respectively. All ROI masks are bilateral. (**A**) Whole-network DMN and MNS masks. (**B**) Within-network ROIs that are thought to be key nodes of either the DMN and MNS. Some of these ROIs are also considered to be nodes of the attentional networks. **Abbreviations:** ROI = region of interest; DMN = default mode network; MNS = mirror-neuron system; mPFC = medial prefrontal cortex; vmPFC = ventromedial prefrontal cortex; PCC = posterior cingulate cortex; TPJ = temporoparietal junction; pIFG = posterior inferior frontal gyrus; dPMC = dorsal premotor cortex; IPS = intraparietal sulcus; LOTC = lateral occipitotemporal cortex; FPCN = frontoparietal control network; VAN = ventral attention network; DAN = dorsal attention network

The masks obtained above were used to extract ROI parameter estimates (mean value of all voxels in a mask) from the single-subject/session Why-How contrasts in the present study. Multiple comparisons across ROIs were accounted for by controlling the false discovery rate (FDR) < 0.05 (106), and *p*-values were adjusted accordingly (*p*_FDR_) through the procedure in (107). Pre-training group differences in neural response were analyzed in a LMEM with Group, Stimulus, and their interaction as effects; the intercept and Stimulus were nested within Subject. In PTSD patients, the relationship between symptom severity (CAPS) and neural response was examined in a LMEM with CAPS and CAPS x Session x Stimulus effects; the intercept, Session, Stimulus, and Session x Stimulus were nested within Subject. The relationship between training outcome (Post-Pre CAPS score difference; CAPSdiff) and neural response was analyzed in a LMEM with CAPSdiff and CAPSdiff x Stimulus as effects; the intercept and Stimulus were nested within Subject.

#### Whole-brain fMRI Analyses

Whole-brain group fMRI analyses were performed to complement the primary ROI analyses. Whole-brain group differences (pre-training) were examined using Aaron Schultz’s MR Tools (108). We specified a general linear model with Group, Stimulus, and their interaction as effects, with the intercept and Stimulus nested within Subject. Residuals from this model were used in AFNI’s 3dFWHMx and 3dClustSim (109) to estimate a cluster extent (*k*) that controls familywise error rate (FWER) < 0.05. *Post-hoc* simple effects tests were conducted in clusters with a significant interaction effect (*p* < 0.005, *k* > 120 voxels).

## Results

### Behavioral Results

The PTSD group featured greater symptom severity (CAPS score) than controls, and symptom severity was marginally reduced after affect labeling training. Full clinical results will be presented in a separate manuscript (94).

Unexpectedly, Why/How task performance (pre-training) did not differ significantly across PTSD and controls. For both response time and accuracy, the main effect of Group and all Group-related interaction effects were not significant (Supplemental Table S1).

### Pre-training Neural Responses across the PTSD and Control Groups

Overall, the Why-How contrast produced activations in DMN ROIs (Figure 3A) and deactivations in MNS ROIs (Figure 3B), aligning with previous studies (87). Unexpectedly, the main effect of Group was not significant in any ROI. Instead, the Group x Stimulus interaction was significant in 4/5 DMN ROIs and 4/5 MNS ROIs. Within these ROIs, *post-hoc* tests revealed that only emotional expressions, not intentional actions, elicited significant Group differences. Specifically, Emotions evoked greater activation in the PTSD group relative to controls (Table 1).

**Figure 3.**
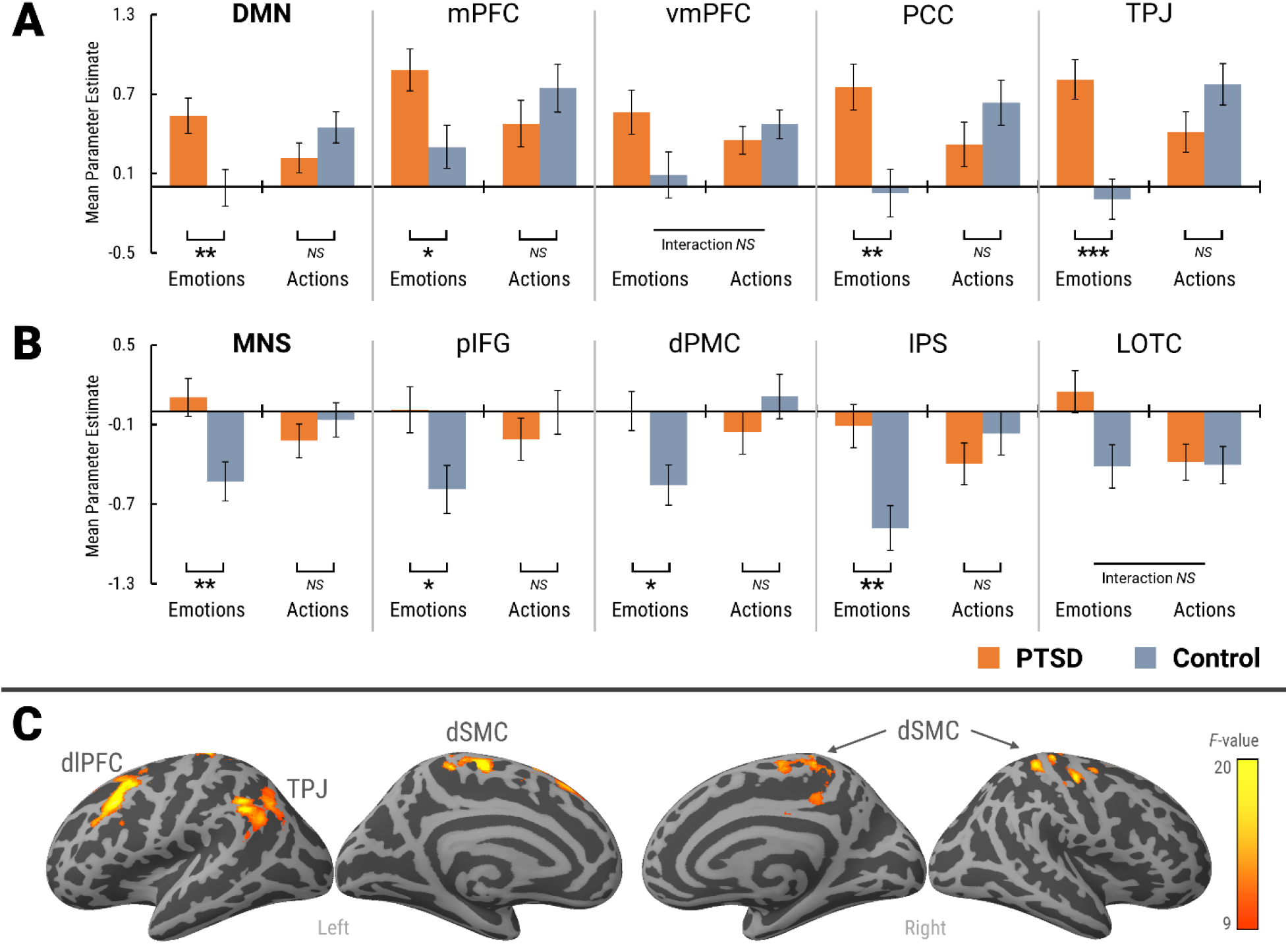
Pre-training Why-How neural responses featured a Group x Stimulus interaction. (**A, B**) Mean parameter estimates across Group and Stimulus in all (**A**) DMN and (**B**) MNS ROIs. Error bars represent standard error of the mean. U-shaped brackets indicate the significance of Group simple effects. (**C**) Whole-brain analysis of the Group x Stimulus interaction (*p* < 0.005, clusterwise FWER < 0.05). Abbreviations: dlPFC = dorsolateral prefrontal cortex; dSMC = dorsal somatomotor cortices; FWER = familywise error rate; \**p*_FDR_ < 0.05; \*\**p*_FDR_ < 0.01; \*\*\**p*_FDR_ < 0.001; ^*NS*^ *p*_FDR_ > 0.05.

**Table 1.** Pre-training Why-How neural responses across Group and Stimulus

| Effect | ROI | DMN ROIs |  |  |  | ROI | MNS ROIs |  |  |  |
| --- | --- | --- | --- | --- | --- | --- | --- | --- | --- | --- |
|  |  | *b* | SE | *t* | *p*<sub>FDR</sub> |  | *b* | SE | *t* | *p*<sub>FDR</sub> |
| Group<sub>PTSD-Control</sub> | DMN | 0.079 | 0.052 | 1.505 | 0.238 | MNS | 0.118 | 0.062 | 1.897 | 0.155 |
| Stimulus<sub>Emotions-Actions</sub> |  | -0.034 | 0.072 | -0.476 | 0.716 |  | -0.035 | 0.074 | -0.464 | 0.716 |
| Group x Stimulus |  | 0.194 | 0.072 | 2.715 | **0.014** |  | 0.198 | 0.074 | 2.656 | **0.014** |
| Group (Emotions)<sup>†</sup> |  | 0.546 | 0.191 | 2.861 | **0.011** |  | 0.633 | 0.207 | 3.054 | **0.009** |
| Group (Actions)<sup>†</sup> |  | -0.232 | 0.162 | -1.426 | 0.387 |  | -0.159 | 0.181 | -0.878 | 0.387 |
| Group<sub>PTSD-Control</sub> | mPFC | 0.079 | 0.098 | 0.808 | 0.422 | pIFG | 0.099 | 0.076 | 1.297 | 0.249 |
| Stimulus<sub>Emotions-Actions</sub> |  | -0.010 | 0.069 | -0.138 | 0.891 |  | -0.090 | 0.093 | -0.970 | 0.671 |
| Group x Stimulus |  | 0.213 | 0.069 | 3.073 | **0.010** |  | 0.200 | 0.093 | 2.162 | **0.043** |
| Group (Emotions)<sup>†</sup> |  | 0.583 | 0.227 | 2.571 | **0.017** |  | 0.599 | 0.251 | 2.383 | **0.023** |
| Group (Actions)<sup>†</sup> |  | -0.268 | 0.251 | -1.065 | 0.387 |  | -0.203 | 0.228 | -0.888 | 0.387 |
| Group<sub>PTSD-Control</sub> | vmPFC | 0.088 | 0.063 | 1.400 | 0.238 | dPMC | 0.071 | 0.083 | 0.862 | 0.422 |
| Stimulus<sub>Emotions-Actions</sub> |  | -0.042 | 0.078 | -0.541 | 0.716 |  | -0.126 | 0.074 | -1.716 | 0.602 |
| Group x Stimulus |  | 0.149 | 0.078 | 1.899 | 0.069 |  | 0.207 | 0.074 | 2.817 | **0.014** |
| Group (Emotions)<sup>†</sup> |  | Interaction NS |  |  |  |  | 0.557 | 0.210 | 2.654 | **0.016** |
| Group (Actions)<sup>†</sup> |  | Interaction NS |  |  |  |  | -0.272 | 0.233 | -1.168 | 0.387 |
| Group<sub>PTSD-Control</sub> | PCC | 0.122 | 0.087 | 1.407 | 0.238 | IPS | 0.135 | 0.070 | 1.919 | 0.155 |
| Stimulus<sub>Emotions-Actions</sub> |  | -0.062 | 0.087 | -0.719 | 0.716 |  | -0.107 | 0.091 | -1.183 | 0.602 |
| Group x Stimulus |  | 0.279 | 0.087 | 3.221 | **0.010** |  | 0.249 | 0.091 | 2.750 | **0.014** |
| Group (Emotions)<sup>†</sup> |  | 0.801 | 0.249 | 3.211 | **0.008** |  | 0.769 | 0.233 | 3.296 | **0.008** |
| Group (Actions)<sup>†</sup> |  | -0.314 | 0.240 | -1.308 | 0.387 |  | -0.228 | 0.226 | -1.010 | 0.387 |
| Group<sub>PTSD-Control</sub> | TPJ | 0.136 | 0.059 | 2.302 | 0.148 | LOTc | 0.146 | 0.066 | 2.223 | 0.148 |
| Stimulus<sub>Emotions-Actions</sub> |  | -0.118 | 0.090 | -1.301 | 0.602 |  | 0.130 | 0.083 | 1.554 | 0.602 |
| Group x Stimulus |  | 0.315 | 0.090 | 3.488 | **0.009** |  | 0.134 | 0.083 | 1.613 | 0.111 |
| Group (Emotions)<sup>†</sup> |  | 0.902 | 0.213 | 4.238 | **0.001** |  | Interaction NS |  |  |  |
| Group (Actions)<sup>†</sup> |  | -0.359 | 0.219 | -1.638 | 0.387 |  | Interaction NS |  |  |  |

<sup>†</sup> *post-hoc* simple effects test; *p*<sub>FDR</sub> = *p*-value adjusted for false discovery rate < 0.05; NS = not significant (*p*<sub>FDR</sub> > 0.05)

Mirroring the ROI results, the whole-brain analysis (Figure 3C) did not find the main effects of Group and Stimulus to be significant (*p* < 0.005, clusterwise FWER < 0.05). Instead, Group x Stimulus was significant in 3 clusters (peak coordinates listed): left dorsolateral prefrontal cortex (dlPFC; x = −33, y = 28, z = 40, *F*_1,33_ = 18.96, *k* = 503), bilateral dorsal somatomotor cortices (dSMC; x = −6, y = −28, z = 60, *F*_1,33_ = 16.73, *k* = 811), and left temporoparietal junction (TPJ; x = −51, y = −51, z = 39, *F*_1,33_ = 16.40, *k* = 476). In all three clusters, *post-hoc* tests revealed that only emotional expressions, not intentional actions, elicited significant Group differences. Specifically, emotional expressions evoked greater activation in PTSD relative to controls in left dlPFC (Emotions: *t*_33_ = 3.60, *p* < 0.001; Actions: *t*_33_ = −1.89, *p* = 0.07), dSMC (Emotions: *t*_33_ = 3.35, *p* = 0.002; Actions: *t*_33_ = −1.94, *p* = 0.06), and left TPJ (Emotions: *t*_33_ = 4.04, *p* < 0.001; Actions: *t*_33_ = −1.98, *p* = 0.06).

### Relationship between Symptom Severity and Neural Responses (PTSD only)

Figure 4 and Table 2 show the relationship between symptom severity and Why-How neural responses in PTSD patients. The main effect of CAPS score was not significant in any ROI. Instead, the CAPS x Session x Stimulus interaction was significant in 4/5 DMN ROIs and 4/5 MNS ROI. *Post-hoc* CAPS simple effects tests revealed that the pattern of interaction was consistent across these ROIs. For emotional stimuli, the CAPS correlation was positive during pre-training (significant in PCC and IPS), and negative during post-training (significant in pIFG). For action stimuli, the CAPS correlation was negative during pre-training (not significant in any ROI), and positive during post-training (significant in DMN, MNS, and IPS).

**Figure 4.**
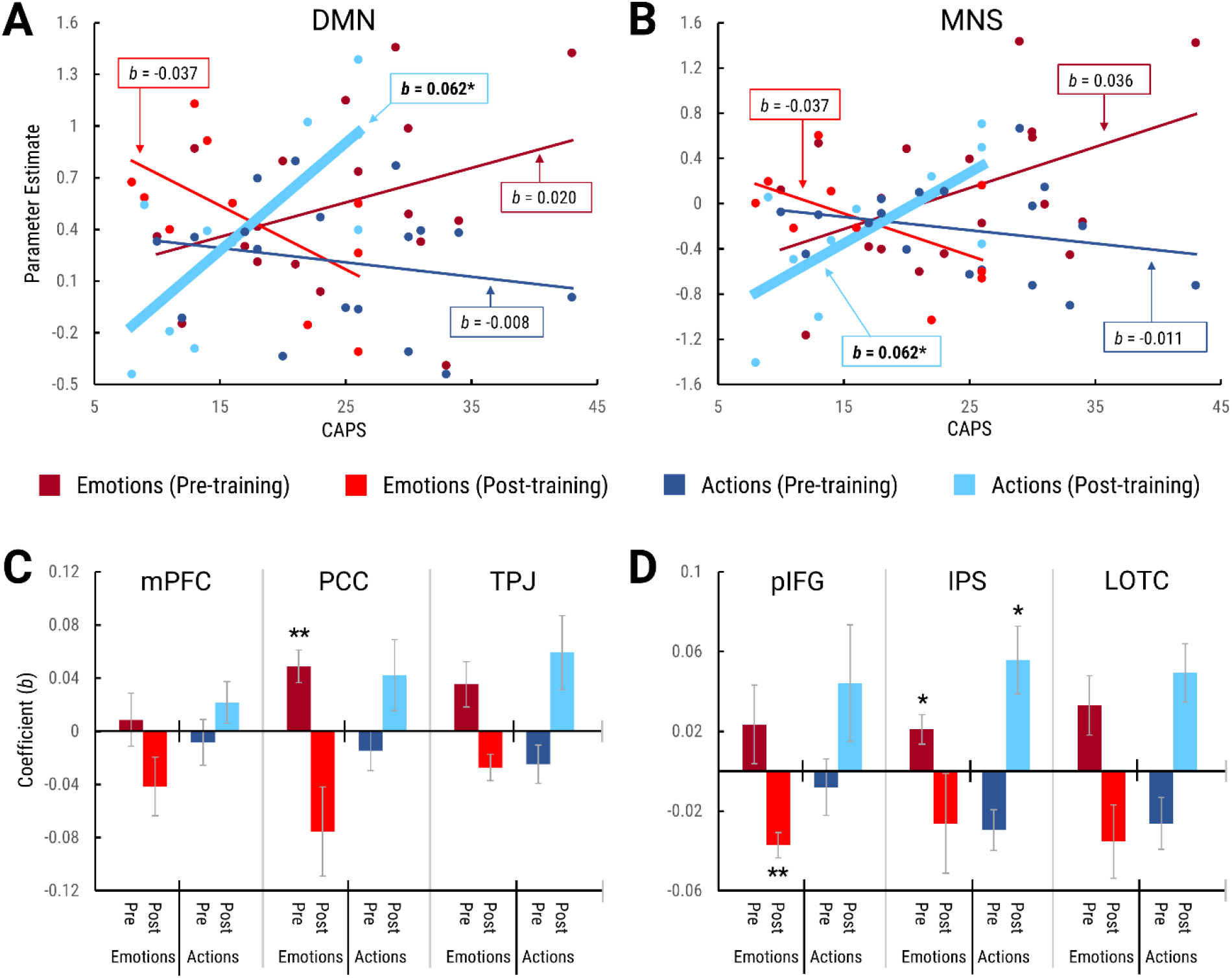
Relationship between symptom severity (CAPS) and Why-How neural responses in PTSD patients. Only ROIs with a significant CAPS x Session x Stimulus interaction are shown. The top panels show scatterplots of parameter estimates and CAPS scores in the **(A)** DMN and **(B)** MNS whole-network masks, with regression lines for CAPS simple effects. Thick lines represent significant regression coefficients, while thin lines represent non-significant regression coefficients. The bottom panels show CAPS simple effects regression coefficients for ROIs within **(C)** DMN and **(D)** MNS. Error bars represent standard error of regression coefficients. **Abbreviations:** CAPS = Clinician-Administered PTSD Scale

**Table 2.**
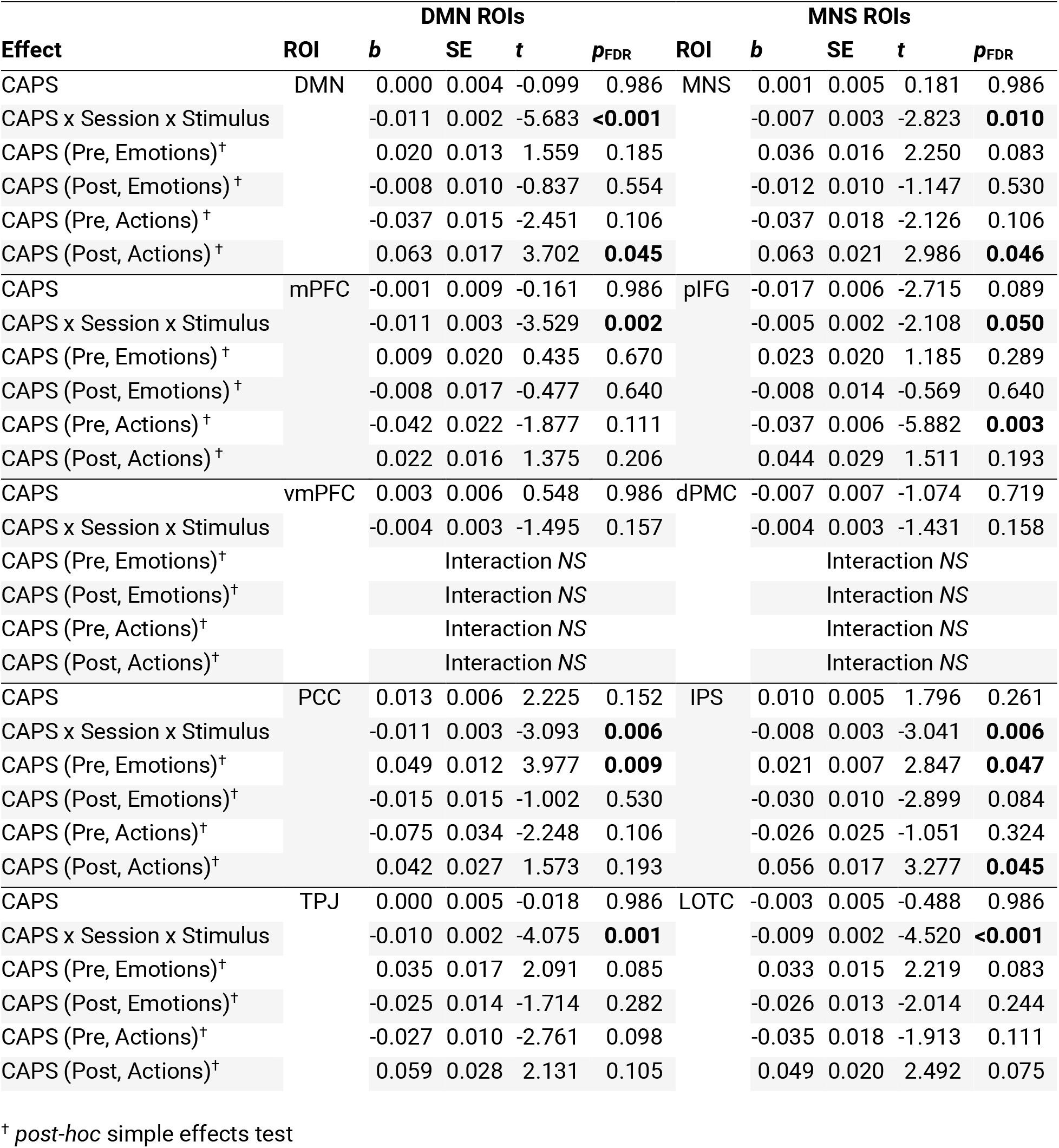
Relationship between symptom severity (CAPS) and Why-How neural activation in PTSD patients

| Effect | ROI | DMN ROIs |  |  |  | ROI | MNS ROIs |  |  |  |
| --- | --- | --- | --- | --- | --- | --- | --- | --- | --- | --- |
|  |  | *b* | SE | *t* | *p*<sub>FDR</sub> |  | *b* | SE | *t* | *p*<sub>FDR</sub> |
| CAPS | DMN | 0.000 | 0.004 | -0.099 | 0.986 | MNS | 0.001 | 0.005 | 0.181 | 0.986 |
| CAPS x Session x Stimulus |  | -0.011 | 0.002 | -5.683 | **<0.001** |  | -0.007 | 0.003 | -2.823 | **0.010** |
| CAPS (Pre, Emotions)<sup>†</sup> |  | 0.020 | 0.013 | 1.559 | 0.185 |  | 0.036 | 0.016 | 2.250 | 0.083 |
| CAPS (Post, Emotions)<sup>†</sup> |  | -0.008 | 0.010 | -0.837 | 0.554 |  | -0.012 | 0.010 | -1.147 | 0.530 |
| CAPS (Pre, Actions)<sup>†</sup> |  | -0.037 | 0.015 | -2.451 | 0.106 |  | -0.037 | 0.018 | -2.126 | 0.106 |
| CAPS (Post, Actions)<sup>†</sup> |  | 0.063 | 0.017 | 3.702 | **0.045** |  | 0.063 | 0.021 | 2.986 | **0.046** |
| CAPS | mPFC | -0.001 | 0.009 | -0.161 | 0.986 | pIFG | -0.017 | 0.006 | -2.715 | 0.089 |
| CAPS x Session x Stimulus |  | -0.011 | 0.003 | -3.529 | **0.002** |  | -0.005 | 0.002 | -2.108 | **0.050** |
| CAPS (Pre, Emotions)<sup>†</sup> |  | 0.009 | 0.020 | 0.435 | 0.670 |  | 0.023 | 0.020 | 1.185 | 0.289 |
| CAPS (Post, Emotions)<sup>†</sup> |  | -0.008 | 0.017 | -0.477 | 0.640 |  | -0.008 | 0.014 | -0.569 | 0.640 |
| CAPS (Pre, Actions)<sup>†</sup> |  | -0.042 | 0.022 | -1.877 | 0.111 |  | -0.037 | 0.006 | -5.882 | **0.003** |
| CAPS (Post, Actions)<sup>†</sup> |  | 0.022 | 0.016 | 1.375 | 0.206 |  | 0.044 | 0.029 | 1.511 | 0.193 |
| CAPS | vmPFC | 0.003 | 0.006 | 0.548 | 0.986 | dPMC | -0.007 | 0.007 | -1.074 | 0.719 |
| CAPS x Session x Stimulus |  | -0.004 | 0.003 | -1.495 | 0.157 |  | -0.004 | 0.003 | -1.431 | 0.158 |
| CAPS (Pre, Emotions)<sup>†</sup> |  | Interaction NS |  |  |  |  | Interaction NS |  |  |  |
| CAPS (Post, Emotions)<sup>†</sup> |  | Interaction NS |  |  |  |  | Interaction NS |  |  |  |
| CAPS (Pre, Actions)<sup>†</sup> |  | Interaction NS |  |  |  |  | Interaction NS |  |  |  |
| CAPS (Post, Actions)<sup>†</sup> |  | Interaction NS |  |  |  |  | Interaction NS |  |  |  |
| CAPS | PCC | 0.013 | 0.006 | 2.225 | 0.152 | IPS | 0.010 | 0.005 | 1.796 | 0.261 |
| CAPS x Session x Stimulus |  | -0.011 | 0.003 | -3.093 | **0.006** |  | -0.008 | 0.003 | -3.041 | **0.006** |
| CAPS (Pre, Emotions)<sup>†</sup> |  | 0.049 | 0.012 | 3.977 | **0.009** |  | 0.021 | 0.007 | 2.847 | **0.047** |
| CAPS (Post, Emotions)<sup>†</sup> |  | -0.015 | 0.015 | -1.002 | 0.530 |  | -0.030 | 0.010 | -2.899 | 0.084 |
| CAPS (Pre, Actions)<sup>†</sup> |  | -0.075 | 0.034 | -2.248 | 0.106 |  | -0.026 | 0.025 | -1.051 | 0.324 |
| CAPS (Post, Actions)<sup>†</sup> |  | 0.042 | 0.027 | 1.573 | 0.193 |  | 0.056 | 0.017 | 3.277 | **0.045** |
| CAPS | TPJ | 0.000 | 0.005 | -0.018 | 0.986 | LOTc | -0.003 | 0.005 | -0.488 | 0.986 |
| CAPS x Session x Stimulus |  | -0.010 | 0.002 | -4.075 | **0.001** |  | -0.009 | 0.002 | -4.520 | **<0.001** |
| CAPS (Pre, Emotions)<sup>†</sup> |  | 0.035 | 0.017 | 2.091 | 0.085 |  | 0.033 | 0.015 | 2.219 | 0.083 |
| CAPS (Post, Emotions)<sup>†</sup> |  | -0.025 | 0.014 | -1.714 | 0.282 |  | -0.026 | 0.013 | -2.014 | 0.244 |
| CAPS (Pre, Actions)<sup>†</sup> |  | -0.027 | 0.010 | -2.761 | 0.098 |  | -0.035 | 0.018 | -1.913 | 0.111 |
| CAPS (Post, Actions)<sup>†</sup> |  | 0.059 | 0.028 | 2.131 | 0.105 |  | 0.049 | 0.020 | 2.492 | 0.075 |

<sup>†</sup> *post-hoc* simple effects test

### Predicting Training Outcomes from Pre-Training Neural Responses (PTSD only)

Figure 5 and Table 3 show the relationship between training outcomes and pre-training Why-How neural responses for PTSD participants who completed affect labeling training. The main effect of Post-Pre CAPS score difference (CAPSdiff) was not significant in any ROI. Instead, CAPSdiff x Stimulus was significant in 3/5 DMN ROIs and 5/5 MNS ROIs. *Post-hoc* CAPSdiff simple effects tests revealed that the pattern of interaction was consistent across these ROIs. The CAPSdiff correlation was negative for emotional stimuli (significant in DMN, PCC, TPJ, MNS, IPS, and LOTC) and flat for action stimuli (not significant in any ROI). In sum, larger neural responses to emotional stimuli predicted larger decreases in symptom severity after affect labeling training.

**Figure 5.**
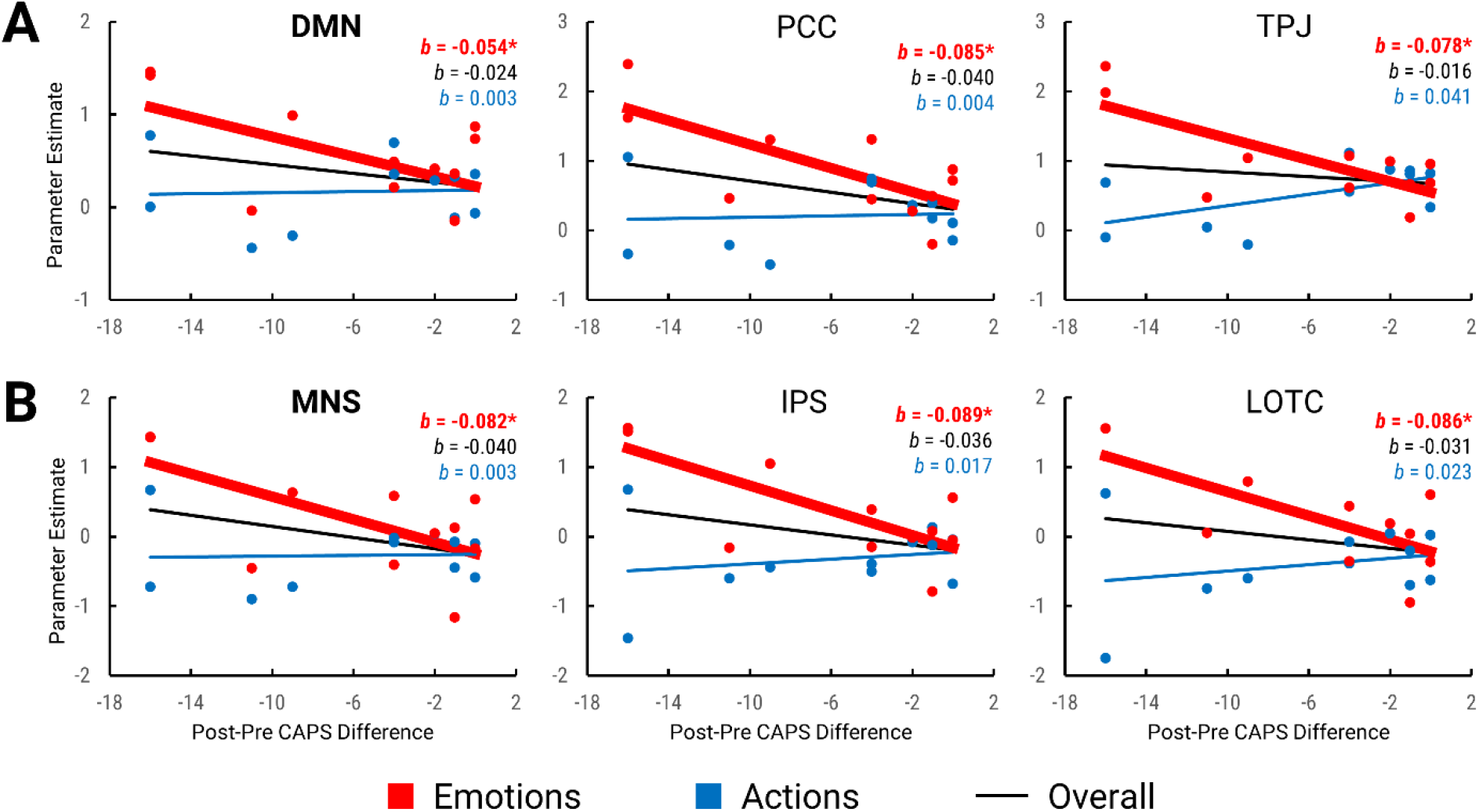
Prediction of training outcomes from pre-training Why-How neural responses in PTSD patients who completed affect labeling training. (**A**) DMN and (**B**) MNS ROIs with significant predictive effects are shown. Scatterplots compare pre-training parameter estimates and Post-Pre CAPS differences, with regression lines plotted for main (overall) and simple (stimulus-specific) effects. Bolded lines represent significant regression coefficients, while thin lines represent non-significant regression coefficients.

**Table 3.**
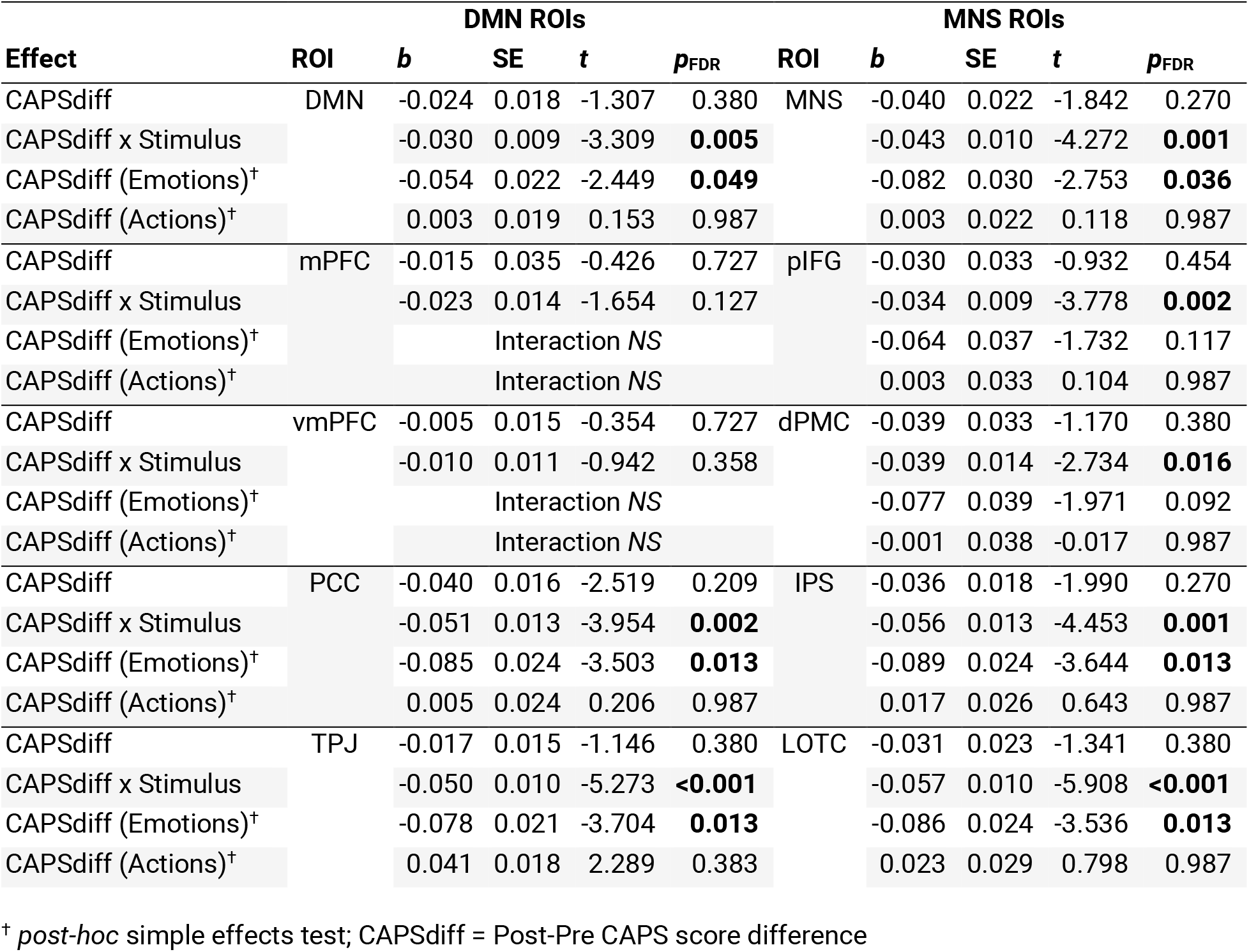
Prediction of training outcomes (CAPSdiff) from pre-training Why-How neural responses in PTSD patients who completed affect labeling therapy.

| Effect | DMN ROIs |  |  |  |  | MNS ROIs |  |  |  |  |
| --- | --- | --- | --- | --- | --- | --- | --- | --- | --- | --- |
|  | ROI | *b* | SE | *t* | *p*<sub>FDR</sub> | ROI | *b* | SE | *t* | *p*<sub>FDR</sub> |
| CAPSdiff | DMN | -0.024 | 0.018 | -1.307 | 0.380 | MNS | -0.040 | 0.022 | -1.842 | 0.270 |
| CAPSdiff x Stimulus |  | -0.030 | 0.009 | -3.309 | **0.005** |  | -0.043 | 0.010 | -4.272 | **0.001** |
| CAPSdiff (Emotions)<sup>†</sup> |  | -0.054 | 0.022 | -2.449 | **0.049** |  | -0.082 | 0.030 | -2.753 | **0.036** |
| CAPSdiff (Actions)<sup>†</sup> |  | 0.003 | 0.019 | 0.153 | 0.987 |  | 0.003 | 0.022 | 0.118 | 0.987 |
| CAPSdiff | mPFC | -0.015 | 0.035 | -0.426 | 0.727 | pIFG | -0.030 | 0.033 | -0.932 | 0.454 |
| CAPSdiff x Stimulus |  | -0.023 | 0.014 | -1.654 | 0.127 |  | -0.034 | 0.009 | -3.778 | **0.002** |
| CAPSdiff (Emotions)<sup>†</sup> |  | Interaction NS |  |  |  |  | -0.064 | 0.037 | -1.732 | 0.117 |
| CAPSdiff (Actions)<sup>†</sup> |  | Interaction NS |  |  |  |  | 0.003 | 0.033 | 0.104 | 0.987 |
| CAPSdiff | vmPFC | -0.005 | 0.015 | -0.354 | 0.727 | dPMC | -0.039 | 0.033 | -1.170 | 0.380 |
| CAPSdiff x Stimulus |  | -0.010 | 0.011 | -0.942 | 0.358 |  | -0.039 | 0.014 | -2.734 | **0.016** |
| CAPSdiff (Emotions)<sup>†</sup> |  | Interaction NS |  |  |  |  | -0.077 | 0.039 | -1.971 | 0.092 |
| CAPSdiff (Actions)<sup>†</sup> |  | Interaction NS |  |  |  |  | -0.001 | 0.038 | -0.017 | 0.987 |
| CAPSdiff | PCC | -0.040 | 0.016 | -2.519 | 0.209 | IPS | -0.036 | 0.018 | -1.990 | 0.270 |
| CAPSdiff x Stimulus |  | -0.051 | 0.013 | -3.954 | **0.002** |  | -0.056 | 0.013 | -4.453 | **0.001** |
| CAPSdiff (Emotions)<sup>†</sup> |  | -0.085 | 0.024 | -3.503 | **0.013** |  | -0.089 | 0.024 | -3.644 | **0.013** |
| CAPSdiff (Actions)<sup>†</sup> |  | 0.005 | 0.024 | 0.206 | 0.987 |  | 0.017 | 0.026 | 0.643 | 0.987 |
| CAPSdiff | TPJ | -0.017 | 0.015 | -1.146 | 0.380 | LOTc | -0.031 | 0.023 | -1.341 | 0.380 |
| CAPSdiff x Stimulus |  | -0.050 | 0.010 | -5.273 | **<0.001** |  | -0.057 | 0.010 | -5.908 | **<0.001** |
| CAPSdiff (Emotions)<sup>†</sup> |  | -0.078 | 0.021 | -3.704 | **0.013** |  | -0.086 | 0.024 | -3.536 | **0.013** |
| CAPSdiff (Actions)<sup>†</sup> |  | 0.041 | 0.018 | 2.289 | 0.383 |  | 0.023 | 0.029 | 0.798 | 0.987 |

<sup>†</sup> *post-hoc* simple effects test; CAPSdiff = Post-Pre CAPS score difference

## Discussion

We conducted the first neuroimaging investigation of social inference in PTSD to help uncover the etiology of PTSD-related social cognitive impairments. To this end, we examined neural activation evoked by the Why/How social inference task (87), which dissociates the two primary networks of the “social brain”: the default mode network (DMN) and mirror neuron system (MNS) (25, 51). We found that DMN and MNS responses differentiated PTSD patients from controls, correlated with symptom severity, and predicted training outcomes. Unexpectedly, these effects were driven almost exclusively by hyperactivation to emotional stimuli. Our neuroimaging results were not corroborated by differences in Why/How task performance, despite numerous reports of impaired social inference performance in PTSD (18). This discrepancy may be attributable to the ease of the task. Taken together, these results suggest that the social brain may be inordinately selective for affective stimuli in PTSD, even in the absence of measurable behavioral impairments.

### Affect-Related Disruption of Social Inference Processing in PTSD

To reveal PTSD-related disruptions in social inference processing, Why-How neural responses were analyzed for PTSD-control differences, symptom severity correlations, and prediction of training outcomes. In all three analyses, emotional facial expressions (Emotions) elicited much stronger PTSD-related effects than intentional hand actions (Actions). In the pretraining session, the PTSD group showed greater activation for Emotions, while controls showed greater activation for Actions. Critically, group differences were only significant for Emotions (Figure 3, Table 1). Similarly, pre-training symptom severity was positively correlated with Emotions-evoked activation and negatively correlated with Actions-evoked activation; these correlations were only significant for Emotions (Figure 4, Table 2). Lastly, better training outcomes were predicted by greater Emotions-evoked activation in the pre-training session (Figure 5, Table 3). These results were generally consistent throughout DMN and MNS, suggesting that both mentalizing and action identification processes are broadly selective for affective stimuli in PTSD. Taken together, affect-selective hyperactivation may be a defining characteristic of social inference processing in PTSD. This aligns with numerous reports of affect-selective hyperactivation in PTSD (68, 81–83, 86, 110).

Given the overarching role of affect in our results, it would be reasonable to expect the strongest effects in ventromedial prefrontal cortex (vmPFC; Figure 2B), the hub of affective processing in the social brain (21, 111–113). Through a wide array of paradigms, vmPFC has been shown to compute the affective valence and value of social and non-social stimuli (114–119). During social inference, vmPFC can represent the emotions of others (40, 120–122). Unexpectedly, vmPFC did not show any significant PTSD-related effects in the current study, though the directionality of effects were consistent with other ROIs. Moreover, whole-brain analyses did not reveal significant PTSD-related effects in core affective regions such as amygdala (123), orbitofrontal cortex (124), and insula (125). Instead, all other ROIs featured significant PTSD-related effects, even though they are less implicated in affective processing (126). Thus, core affective processes may not play a key role in disrupting social inference processing in PTSD.

Alternatively, the affect-selective hyperactivation we observed may not reflect altered affect *per se*, but rather selective processing of affective information by the wide array of social cognitive processes subserved by DMN and MNS. Indeed, PTSD-related effects were stronger in the whole-network DMN and MNS masks, indicating that social inference processing was disrupted on a network-wide basis. Moreover, the within-network ROIs featured similar patterns of results as the whole-network masks, though effects were often weaker or not significant. Nevertheless, the consistency of these effects is remarkable given the functional heterogeneity between and within DMN and MNS (122, 127).

### An Attentional Account of Social Inference Dysfunction in PTSD

Outside of core affective processes, what neurocognitive mechanisms could instigate such broad affect-selectivity throughout the social brain? Putatively, attention may be one such mechanism, as attentional processes are frequently reported to be inordinately biased towards emotional stimuli in PTSD (82, 77, 128–136). Concordantly, PTSD-related attentional biases have been linked with affect-evoked hyperactivation throughout DMN (80, 81). Moreover, DMN activation has been shown to correspond with attention level during social tasks (48, 88, 137–139). Similarly, MNS activation appears to be modulated by both top-down and bottom-up attention (41, 88, 140–144).

An attentional account is further supported by the anatomical overlap between the attention networks and regions with significant PTSD-related effects in the present study. With the exception of medial prefrontal cortex, ROIs with significant effects appear to overlap with either the dorsal attention (DAN), ventral attention (VAN), or frontoparietal control (FPCN) networks. DAN is involved in top-down attention, and includes intraparietal sulcus (IPS), dorsal premotor cortex (dPMC), and lateral occipitotemporal cortex (LOTC) (58, 145, 146). VAN, involved in bottom-up attention and attentional reorientation, includes temporoparietal junction (TPJ) and posterior inferior frontal gyrus (pIFG) (74, 146). FPCN includes parts of posterior cingulate cortex (PCC), and is thought to facilitate attentional control by mediating activity between DMN, DAN, and other networks (147–150). Moreover, whole-brain analyses revealed PTSD-related effects in one region outside our *a priori* ROIs: the dorsolateral prefrontal cortex, a central node of DAN and FPCN (145, 151). Accordingly, other studies have observed affect-evoked hyperactivation in DAN, VAN, and FPCN in PTSD (76, 80–83, 85). Taken together, affective attentional biases in PTSD may drive widespread affect-selective hyperactivation throughout DMN and MNS during social inference.

### Affect Labeling Training

The PTSD group underwent affect labeling training, which involves labeling the emotional content of stimuli (90). Affect labeling is an emotional inhibitory regulation strategy that has been found to downregulate amygdala responses via right ventrolateral prefrontal cortex (vlPFC) in healthy subjects (90, 91, 152, 153). Though we did not find PTSD-related effects in amygdala or vlPFC, affect labeling training was found to reduce symptom severity (94). Affect labeling may inhibit the affective components of social inference processing, as reactivity to emotional stimuli became negatively correlated with symptom severity after training—a reversal of the positive correlation found prior to training (Figure 4). This posttraining negative correlation reached significance only in pIFG, perhaps signifying the importance of mirroring and bottom-up attentional processes during socioaffective inhibitory regulation (32, 146). Moreover, better training outcomes were predicted by higher Emotions-evoked activation in the pre-training session (Figure 5), suggesting that engagement with emotional stimuli enhances the efficacy of affect labeling training—an interpretation consistent with other studies on exposure-based PTSD interventions (154–157). In sum, affect labeling training may be better suited for patients with greater affect-selective hyperactivation during social inference.

### Limitations and Future Directions

The interpretation of these results should be tempered by the relatively small sample size of this study, especially in the post-training analyses. Additionally, generalizability may be limited by our selective recruitment of American veterans exposed to combat trauma. Future studies should use larger and more diverse samples. A potential confound in this study are the non-affective stimulus differences between Emotions (faces) and Actions (hands); our key finding of Emotions-selective hyperactivation may not be exclusively driven by affect. Future studies should better match emotional and non-emotional stimuli. Another caveat is the putative nature of the functional-anatomic overlap between our findings and the attention networks. This overlap was inferred using reverse inference from existing literature—a form of reasoning that can be tenuous (158, 159). This functional-anatomic overlap could be more definitively investigated by including functional localizers for DAN, VAN, and FPCN in addition to DMN and MNS. More broadly, techniques such as multivariate analyses, connectivity analyses, and other neuroimaging modalities would be useful in further characterizing PTSD-related neural dynamics during social inference.

### Conclusion

In the first neuroimaging investigation of social inference in PTSD, the social brain was found to be broadly selective towards emotional stimuli in PTSD. Affect-selective hyperactivation throughout DMN, MNS, and beyond differentiated PTSD patients from controls, correlated with symptom severity, and predicted training outcomes. Despite this, PTSD-related effects were not significant in core affective regions. Instead, our data putatively highlight the role of attentional processes in disrupting social inference processing in PTSD. These results indicate that further study of social inference processing in PTSD is strongly warranted, specifically in disentangling the roles of affect and attention.

## Acknowledgements

This research was supported by the Defense Advanced Research Projects Agency (DARPA), an agency of the U.S. Department of Defense, through the Army Research Laboratory (PIs: Burklund, Craske & Lieberman; W911NF-14-C-0056). The author Kevin M. Tan was supported by the National Science Foundation Graduate Research Fellowship Program (DGE-1650604).

