## Supplemental Table 1 for "Posttraumatic Stress Disorder and the Social Brain: Affect-Related Disruption of the Default and Mirror Networks"

**Response Time (ms)**

| Effect | *b* | SE | *t* | *p* |
|---|---|---|---|---|
| Intercept (grand mean) | 594.12 | 21.94 | 27.082 | **<0.001** |
| $Group_{PTSD-Control}$ | -16.79 | 21.94 | -0.765 | 0.444 |
| $Prompt_{Why-How}$ | 23.84 | 4.87 | 4.891 | **<0.001** |
| $Stimulus_{Emotions-Actions}$ | -42.81 | 4.64 | -9.218 | **<0.001** |
| Group x Prompt | 5.08 | 4.87 | 1.042 | 0.298 |
| Group x Stimulus | 2.50 | 4.64 | 0.538 | 0.591 |
| Prompt x Stimulus | 12.22 | 4.32 | 2.827 | **0.005** |
| Group x Prompt x Stimulus | -0.85 | 4.32 | -0.197 | 0.844 |

**Accuracy (%)**

| Effect | *b* | SE | *t* | *P* |
|---|---|---|---|---|
| Intercept (grand mean) | 93.73 | 0.84 | 112.220 | **<0.001** |
| $Group_{PTSD-Control}$ | -0.24 | 0.84 | -0.291 | 0.771 |
| $Prompt_{Why-How}$ | -1.22 | 0.38 | -3.222 | **0.001** |
| $Stimulus_{Emotions-Actions}$ | -0.12 | 0.37 | -0.318 | 0.751 |
| Group x Prompt | 0.19 | 0.38 | 0.509 | 0.611 |
| Group x Stimulus | 0.09 | 0.37 | 0.244 | 0.807 |
| Prompt x Stimulus | -0.48 | 0.37 | -1.311 | 0.190 |
| Group x Prompt x Stimulus | 0.31 | 0.37 | 0.850 | 0.395 |

**Supplemental Table S1.** LMEMs of Why/How task performance during the pre-treatment fMRI session. For both response time and accuracy, there were no significant main or interaction effects related to differences between the PTSD and control groups. The only significant effects were related to differences between task conditions. The *Why* prompt elicited significantly slower responses and reduced accuracy relative to the *How* prompt. *Emotions* stimuli elicited significantly faster RTs than *Actions* stimuli. For RTs, there was also a significant interaction effect between Prompt and Stimulus. No other effects were significant. **Abbreviations:** ms = millisecond; LMEM = linear mixed-effect model
